# Amphibian-derived peptide analog TB_KKG6K: A powerful drug candidate against *Candida albicans* with anti-biofilm efficacy

**DOI:** 10.1101/2025.08.22.671737

**Authors:** Cristina Schöpf, Anik Geschwindt, Magdalena Knapp, Anna C. Seybold, Débora C. Coraça-Huber, Michael J. Ausserlechner, Alessandra Romanelli, Florentine Marx

## Abstract

*Candida albicans*, a commensal and opportunistic fungal pathogen, is a major clinical concern due to its ability to cause infections ranging from mild mucosal conditions to life-threatening systemic diseases, particularly in immunocompromised patients. Its capacity to form biofilms on medical devices further complicates treatment by enhancing antifungal resistance and immune evasion. In the search for novel therapeutic strategies, the lysine-enriched amphibian-derived temporin B analog, TB_KKG6K, has emerged as a promising antifungal agent. This study demonstrates that TB_KKG6K exhibits potent activity against planktonic *C. albicans* cells, with a low potential to induce adaptation or resistance, even after prolonged exposure. TB_KKG6K has no adverse impact on the anti-*Candida* efficacy of standard antifungal drugs like amphotericin B, caspofungin, fluconazole or 5-flucytosine, when applied in combination. Additionally, TB_KKG6K effectively reduces biofilm maturation on silicone elastomers, a material commonly used in medical devices, further highlighting its therapeutic potential. These data together with our previous documentation of minimal cytotoxicity and irritation potential in human cells, makes TB_KKG6K a strong candidate for combating both planktonic and biofilm-associated *C. albicans* infections. These findings underscore the dual efficacy of TB_KKG6K and its potential to address the challenges posed by *C. albicans* in clinical settings.

## Introduction

*Candida albicans* is a commensal and opportunistic pathogen commonly found within the human microbiota. It is capable of forming complex biofilms, in which different cell types (yeast-like cells, germinating cells, pseudohyphae and hyphae) are embedded in a protective extracellular matrix (ECM), composed of glycoproteins, polysaccharides, lipids and nucleic acids [1–3]. *C. albicans* biofilms are associated with a range of clinical manifestations, from mild local infections to life-threatening conditions such as candidemia, which may result in high mortality rates in patients with comorbidities [4–6]. The protective biofilm environment substantially enhances the pathogenic potential of *C. albicans* by promoting resistance to antifungal drugs and evasion of host immune defenses, a matter of significant concern within the clinical context [1, 7–9].

Biofilm formation on abiotic surfaces is a major factor in device-associated complication in the clinics where *C. albicans* causes persistent hospital-acquired infections resistant to antifungal treatment, and compromises the functionality of implanted medical devices, necessitating premature replacement and contributing to escalating healthcare costs [10–13]. *C. albicans* is able to form biofilms on almost any type of medical equipment, including systemic and topical devices, such as contact lenses and dentures, and those that come into contact with or traverse the skin, particularly synthetic polymers such as silicone found in catheters or prostheses [1, 8, 14]. Adherence of *C. albicans* to silicone and subsequent biofilm formation has been shown to reduce the efficacy of standard antifungal drugs like azoles, polyenes or echinocandins, due to the development of resistance mechanisms [15–17]. This situation highlights the necessity to search for new antifungal molecules that are effective against yeast biofilms.

Antimicrobial peptides (AMPs) have been identified as promising candidates for the development of novel antimicrobial therapeutic agents. The European red frog *Rana temporaria* secretes the AMPs, temporins, by granular glands to protect their skin from infection with microbial pathogens. These short (8-14 amino acid long), mildly cationic (0 to +3 at pH 7) peptides belong to one of the biggest AMP families in nature [18]. Rational design and chemical modifications enabled the creation of peptide analogs with different primary structures and improved efficacy in comparison to their parent peptides. One promising example is temporin B (TB). The lysine-enriched TB analog TB_KKG6K (KKLLPIVKNLLKSLL; molecular weight [MW], 1,718.2 Da) exhibits an increased antimicrobial spectrum and enhanced tolerability in the host compared to TB [19]. We could recently show that this peptide analog acts in a fungicidal way against planktonic and sessile *C. albicans* cells *in vitro*. Its mode of action affects the cell membrane function, induces the production of intracellular reactive oxygen species (iROS) and results in the disintegration of the subcellular structures in the yeast cells, while showing low cytotoxicity in human primary cells and low irritation potential in three-dimensional (3D) reconstructed human skin [20–22].

In the present study we wanted to investigate in more detail the anti-*Candida* efficacy of TB_KKG6K. By combining microbiological and molecular biology analyses, and high-end microscopy, we collected data which for the first time provide evidence for the low potential of TB_KKG6K to induce adaptation or resistance in planktonic *C. albicans* cells and a comprehensive insight into its efficacy to reduce biofilm matured on the surface of a medically relevant silicone elastomer.

## Material and methods

If not otherwise stated, the chemicals and compounds used in this study were purchased from Sigma-Aldrich, St. Louis, MO, USA. All media and solutions used are summarized in Table 1.

**Table 1.** Media and solutions used in this study.

| Media/Solution | Composition<sup>§</sup> |
| --- | --- |
| Potato Dextrose Broth (PDB) | 0.65% Potato infusion, 2% Glucose |
| Potato Dextrose Agar (PDA) | Potato Dextrose Broth, 2% Agar |
| Phosphate Buffered Saline (PBS) | 0.05% KH<sub>2</sub>PO<sub>4</sub>, 0.28% K<sub>2</sub>HPO<sub>4</sub>, 0.9% NaCl, pH 7.4 |
| Dulbecco's phosphate-buffered saline (D-PBS) | 0.02% KCl, 0.02% KH<sub>2</sub>PO<sub>4</sub>, 0.8% NaCl, 0.115% Na<sub>2</sub>HPO<sub>4</sub>, pH 7.0 |
| Yeast Peptone D-Glucose (YPD) Agar | 1% Yeast extract, 2% Peptone, 2% Glucose, 2% Agar |

<sup>§</sup>Percent values (%) are depicted as (wt/vol).

### Peptide synthesis

TB_KKG6K was synthesized and purified by reversed-phase high performance liquid chromatography (RP-HPLC) as described previously [21].

### C. albicans cultivation

*C. albicans* strain CBS 5982 was grown overnight in 5 mL of PDB (Carl Roth, Karlsruhe, Germany) at 37°C with shaking at 200 rpm.

### Checkerboard assay

The inhibitory concentration of TB_KKG6K and antifungals was defined as the concentration that reduced microbial growth ≥ 90% (IC_90_). The antifungal potential of TB_KKG6K, alone and in combination with amphotericin B, fluconazole (Santa Cruz Biotechnology, Dallas, Tx, USA), caspofungin, and 5-flucytosine (5-FC; TCI Deutschland GmbH, Eschborn, Germany), was evaluated using the checkerboard method based on the broth microdilution technique as previously described [20, 23]. The compounds were tested at concentrations starting at twice their respective IC_90_ values, previously determined by a broth microdilution assay in 0.05x PDB, and then serially diluted to lower concentrations. One antifungal was diluted along the X-axis (columns) and the second along the Y-axis (rows), creating a matrix of unique combinations of drug concentrations. A 50 μL aliquot of each four-fold concentrated antifungal compound, combined with 50 μL of the second four-fold concentrated compound or 50 μL of medium (control for single compound testing), was dispensed into a 96-well microplate and mixed with 100 μL of a 10⁴ cells mL^-1^ inoculum, resulting in a final volume of 200 μL. The microplates were incubated at 30°C for 24 h, and growth was assessed visually and spectrophotometrically. The results were expressed as the percentage of growth relative to the untreated growth control.

Antifungal interactions of the individual drug combinations were analyzed using the fractional inhibitory concentration index (FICI) as described [23, 24]. The FICI was calculated by summing the fractional inhibitory concentrations (FICs) for each compound, based on the IC_90_ values of the individual antifungals (A and B) both in combination and individually: ΣFIC = FIC_A_ + FIC_B_. FIC_A_ refers to the IC_90_ of drug A in combination/IC_90_ of drug A alone and FIC_B_ to the IC_90_ of drug B in combination/IC_90_ of drug B alone. FICI was defined as the lowest ΣFIC determined in three independent experiments. Interactions were classified synergistic (FICI ≤ 0.5), additive/indifferent (0.5 < FICI < 4), and antagonistic (FICI > 4) [23].

### Drug adaptation experiment

A drug adaptation experiment was performed *in vitro* according to Papp et al. 2018 [25] and modified as described in [26]. In brief, three individual colonies of *C. albicans* per condition (treatment with TB_KKG6K and fluconazole, respectively, and untreated growth control) were picked from a PDA plate and grown in 0.05x PDB, resulting in three lineages per condition. Then, a suspension of 3x 10^4^ cells mL^-1^ of *C. albicans* was prepared in 0.05x PDB in triplicate for each lineage. For the growth control, 1 mL of the cell suspension was transferred to 2 mL of drug-free 0.05x PDB medium. For the antifungal treatment, 1 mL of the cell suspension was mixed with 2 mL of 0.05x PDB medium containing half the IC_90_ (0.5x IC_90_) of TB_KKG6K (1 µM) or fluconazole (3.2 µM), which served as the positive control. The cells remained untreated for the positive growth control. All samples were incubated at 30° C for 24 h and with shaking at 200 rpm. Then, 100 µL of each sample was transferred to 3 mL of fresh medium, maintaining the 0.5x IC_90_. This subculturing process was repeated at 24-hour intervals for a total of five days. Subsequently, the cells were diluted 1:30 into medium supplemented with the doubled concentration of antifungal compound. Subculturing was continued in 24 h intervals at this antifungal concentration for three days. Then the concentration of the antifungal compound was doubled again. These cultivation cycles were repeated until no cells proliferated any more or an antifungal compound concentration of 32x IC_90_ was reached. The untreated cells of the growth control were transferred on a daily basis to fresh, but drug-free, 0.05x PDB medium. Throughout the experiment, the growth was monitored spectrophotometrically using a multimode microplate reader (FLUOstar Omega, BMG Labtech, Ortenberg, Germany) by determining the OD_620_ of a 200 µL sample at the end of the third subculturing step of each concentration cycle. When very low OD_620_ values were reached, a 100 µL sample of the culture was streaked out on YPD agar and incubated at 37°C for 24 h to count the surviving cells (colony forming units, CFU).

### *C. albicans* biofilm cultivation on silicone elastomer discs

Discs were laser-cut from silicone sheets (0.25 mm, MVQ Silicones GmbH, Weinheim, Germany) with diameters of 9 mm for 48-well plates (VWR, Randnor, PA, USA) and 14 mm for 24-well plates (CytoOne, Starlab, Hamburg, Germany). They were washed by vortexing in 15 mL of ddH_2_O, followed by sonication (35 kHz; Bandelin Sonorex, BANDELIN electronic GmbH & Co. KG, Berlin, Germany) for 10 min. The water was replaced, and this process was repeated twice. Subsequently, the discs were immersed in 15 mL of 70% ethanol for 24 h. Discs were dried under sterile conditions in a safety cabinet and exposed to UV light on both sides for 30 min each. The discs were stored at room temperature until usage.

Before seeding *C. albicans* cells, the discs were placed in a 48-well or 24-well plate and covered with heat-inactivated, sterile fetal bovine serum (PAN-Biotech GmbH, Aidenbach, Germany). They were gently shaken at 100 rpm for 10 min and then incubated statically at 37°C overnight. The next day, the discs were washed with 0.5-1 mL of sterile PBS for 10 min under gentle shaking (100 rpm). The PBS was removed, and the discs were air-dried under sterile conditions before being transferred to a new plate.

Then, 500 µL from a freshly diluted overnight culture of *C. albicans* containing 10⁶ cells mL^-1^ in 0.05x PDB were seeded in each well of a 48-well plate equipped with 9 mm diameter silicone discs, resulting in a cell density of 5x 10^5^ per well. In 24-well plates equipped with 14 mm diameter silicone discs, the cell density was adjusted to 5x 10⁶ cells mL^-1^. A total volume of 1.2 mL (6x 10⁶ cells) was seeded into each well. Cells were distributed in circular motions and incubated statically at 30°C for up to 72 h. The medium was exchanged every 24 h.

### Antifungal therapy

The anti-*Candida* efficacy of 2-50 µM TB_KKG6K was tested on a 48 h-matured biofilm and compared to controls treated with amphotericin B (1.25 µg mL^-1^). An untreated sample was included as growth control. Samples were analyzed after 4 h or 24 h of incubation at 30°C.

### Viability assay of sessile *C. albicans* cells

Discs were harvested by aspiration of non-adherent cells and placed in a 2 mL reaction tube with 1 mL of sterile PBS. They were vortexed at the highest setting for 30 sec, followed by three cycles of sonication (35 kHz) in a water bath (Bandelin Sonorex) for 1 min each as previously described [27]. Vortexing was then repeated. Serial dilutions were prepared, and 100 µL of each dilution was plated on PDA. Plates were incubated for 24 h at 37°C for the quantification of CFU

### RNA extraction and quantitative PCR

Total RNA was isolated from sessile cells grown on 14 mm silicone discs in 24-well plates. Biofilm that had matured for 48 h was subjected to treatment with 5 µM TB_KKG6K for a period of 4 h. Then, eight discs per condition were pooled for RNA extraction and cells were detached by rigorous pipetting. The disruption of cells was achieved mechanically by employing glass beads (0.50-0.75 mm, RETSCH GmbH, Haan, Germany) in 1 mL of TRI reagent (TRI Reagent®). This process was conducted thrice in a Mixer Mill (MM400, RETSCH GmbH) for 2 min at a frequency of 30 Hz. The tubes were cooled for 1 min on ice between each repetition. Subsequently, 200 μl of chloroform was added to each tube, and the tubes were vortexed for 15 sec and left to incubate at room temperature for 5 min. Then the tubes were centrifuged at 4°C and 12,000 G for 10 min. The upper phase containing the RNA was transferred to a 1.5 mL microcentrifuge tube, and an equal volume of 70% (vol/vol) isopropanol was added. A quantity of 20 µg of glycogen was added to the solution, after which the RNA was precipitated at -20°C overnight. After centrifugation of the samples at 4°C and 12,000 G for 10 min the isopropanol was removed, and the RNA pellet was washed twice in 500 µL of 75% (vol/vol) ethanol. For elution, 20 μl of RNase-free water were used. Subsequently, the RNA was treated with 2 units of DNAse I (RNase free, New England Biolabs, Inc., Ipswich, MA, USA) per 10 µg of RNA at 37°C for 30 min. The RNA was purified by phenol-chloroform-isoamylalcohol (Carl Roth) extraction and the concentration and quality of RNA were analyzed using a NanoPhotometer™ (NP80, Implen GmbH, Munich, Germany).

One μg of RNA was utilized for cDNA synthesis (iScriptTM RT Supermix, Bio-Rad Laboratories, Bio-Rad Laboratories, Hercules, CA, USA). The reaction was performed using the PikoReal 96 System (PikoReal 96 Realtime PCR System, Thermo Fisher Scientific, Waltham, MA, USA). The iQTM SYBR Green Supermix kit (Bio-Rad Laboratories) was applied for qPCR, with a total volume of 20 μL per reaction. An initial evaluation of the primer efficiency of the primers listed in Table 2 was conducted prior to this study. For each reaction, 50 ng of cDNA were used. The gene expression fold change was calculated using the ΔΔ Ct values [28].

**Table 2.**
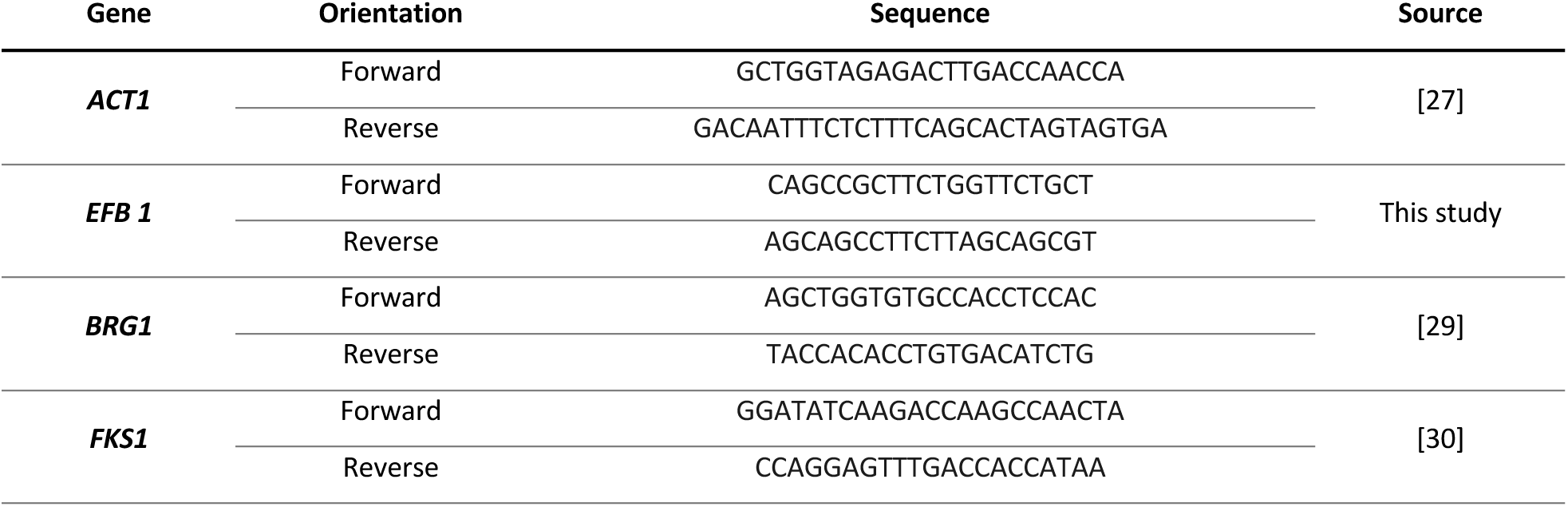
Primers used in this study.

### Scanning electron microscopy (SEM)

Discs were harvested by removing the media and transferring them into a new 48-well plate. They were fixed in 2.5% glutaraldehyde (vol/vol in PBS) at 4°C for 24 h. Dehydration was performed using an ascending ethanol (50-70-80-90%) or acetone series (50-70-80-90-100%). The discs were then mounted on aluminum pins and sputter-coated with gold. Microscopy was performed using JSM-6010LV (JEOL GmbH, Freising, Germany) or TESCAN CLARA SEM (TESCAN GROUP, Brno, Czech Republic).

### Confocal microscopy

Biofilm samples were fixed on the silicone discs in 500 µL of 4% (wt/vol) paraformaldehyde in PBS for 30 min at room temperature. The fixative was then removed, and the discs were washed with 500 µL of PBS for 10 min. Staining was started with Concanavalin A conjugate (100 µg mL^-1^, Alexa Fluor 633 [AF633], Thermo Fisher Scientific) for 30 min. Then, the biofilm was thoroughly washed with D-PBS and subsequently stained with Calcofluor White (1 mg mL^-1^) for 2 min, before washing the samples again with D-PBS to ensure the removal of any residual stain. Subsequently, the biofilm on the silicone discs was mounted in 7 µL of Fluoroshield™ and covered with a 12 mm high precision coverslip (Thorlabs Inc., Newton, NJ, USA). Biofilm was analyzed using a SP8 gSTED microscope (Leica Microsystems GmbH, Wetzlar, Germany). All recordings were processed using the same parameters in Huygens Professional 25.04 (Scientific Volume Imaging B.V., Hilversum, The Netherlands).

### Statistics

Statistical analysis was conducted using Prism 9.1.0 (GraphPad Software, San Diego, CA, USA). Values are given as mean ± standard deviation (SD) per experimental setting (n = 3) and statistical significance (\**P* ≤ 0.05; ** *P* ≤ 0.005) was determined by one-way ANOVA, followed by Dunnett’s test, if not stated otherwise.

## Results

### TB_KKG6K has low potential to induce adaptive mechanisms in *C. albicans*

Our first aim was to evaluate the potential of TB_KKG6K to induce resistance in *C. albicans* over successive generations grown under controlled *in vitro* conditions. Cells treated with fluconazole served as a positive control for adaptation and resistance induction. This fungistatic drug is known to elicit adaptive responses in *C. albicans* and has the potential to induce resistance mechanisms [31–33]. An untreated control served to monitor the growth in the absence of antifungals.

The three lineages of *C. albicans* cells exhibited varying susceptibilities towards 1x IC_90_ after prolonged exposure (120 h) to the subinhibitory concentration (0.5x IC_90_) of TB_KKG6K. However, none of the lineages were able to survive serial passages at concentrations exceeding 1x IC_90_ of TB_KKG6K as the OD_620_ values dropped to zero (Figure 1A). This was confirmed by plating aliquots from the third passage of each lineage exposed to 2x IC_90_ TB_KKG6K on YPD plates without antifungal peptide supplementation. No CFU could be counted after the incubation at 37°C for 24 h. In contrast, cells exposed to gradually increasing concentrations of fluconazole demonstrated the ability to adapt and survive treatment at drug concentration as high as 32x IC_90_. Initially, all three *C. albicans* lineages showed a decrease in OD_620_ values upon exposure to 1x IC_90_ of fluconazole. However, a steady recovery was observed over the time, as indicated by progressively increasing OD_620_ values throughout the cultivation period (Figure 1B). Meanwhile, the untreated lineages of the growth control maintained relatively stable OD_620_ values for the entire duration of the experiment (Figure 1C).

**Figure 1.**
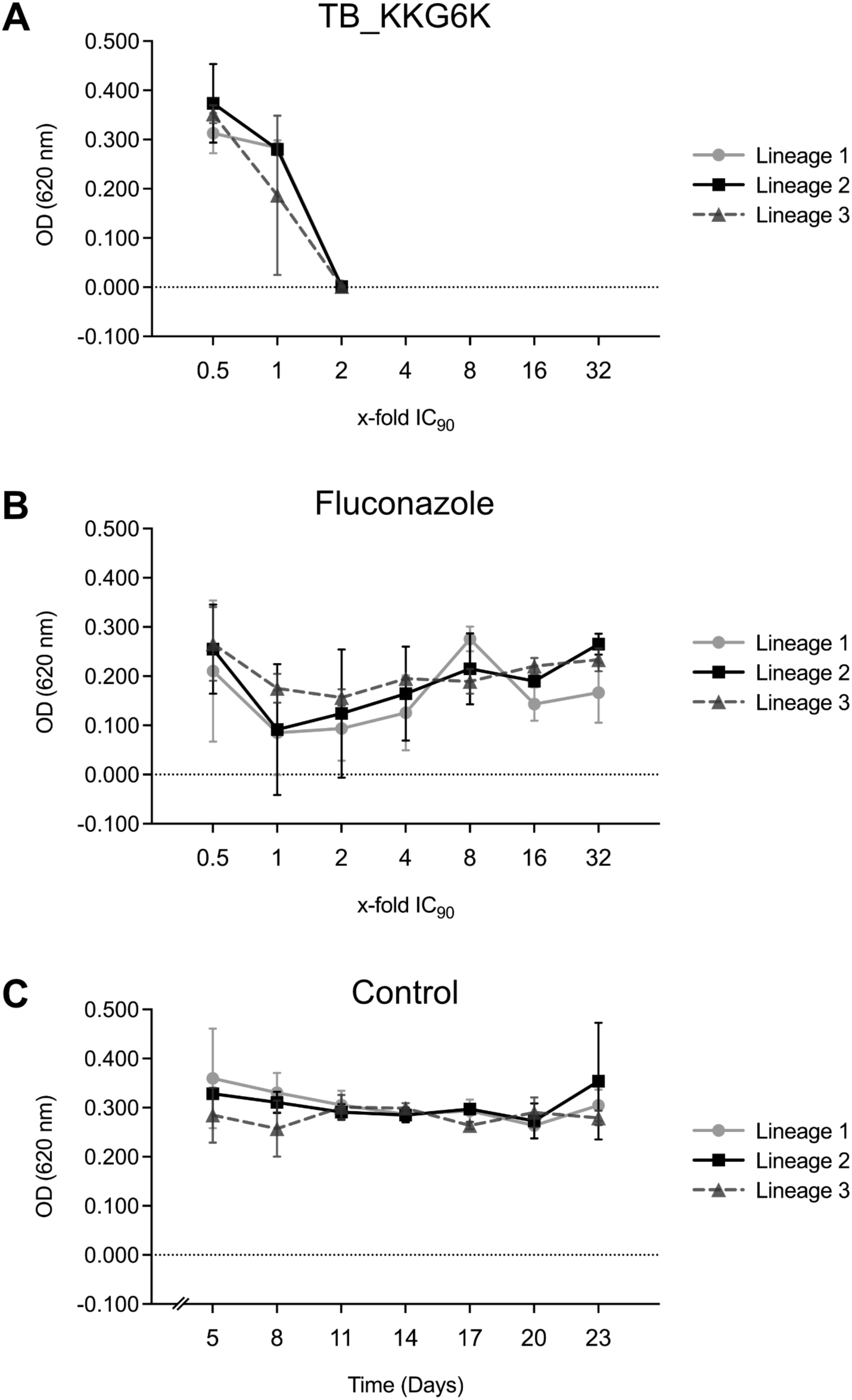
Adaptation of *C. albicans* to increasing concentrations of antifungals. *C. albicans* was exposed to increasing concentrations of (A) TB_KKG6K and (B) fluconazole, or (C) was left untreated (Control). The OD_620_ of three independent lineages, each cultivated in triplicates, were measured at the final passage of the cells in the respective concentrations (in 72 h intervals).

### TB_KKG6K shows no adverse interference with standard antifungal drugs

Next, we investigated the interaction between the peptide TB_KKG6K and one of the four licensed antimycotic drugs on *C. albicans in vitro*. The drugs amphotericin B, caspofungin, fluconazole and 5-FC in combination with the peptide were tested in a checkerboard assay, after having determined the individual IC_90_ values for each antifungal in a standard broth microdilution assay. As summarized in Table 3, TB_KKG6K showed additive/indifferent effects against *C. albicans* with amphotericin B (FICI 0.75), caspofungin (FICI 0.63), fluconazole (FICI 1.5), and 5-FC (FICI 2).

**Table 3.**
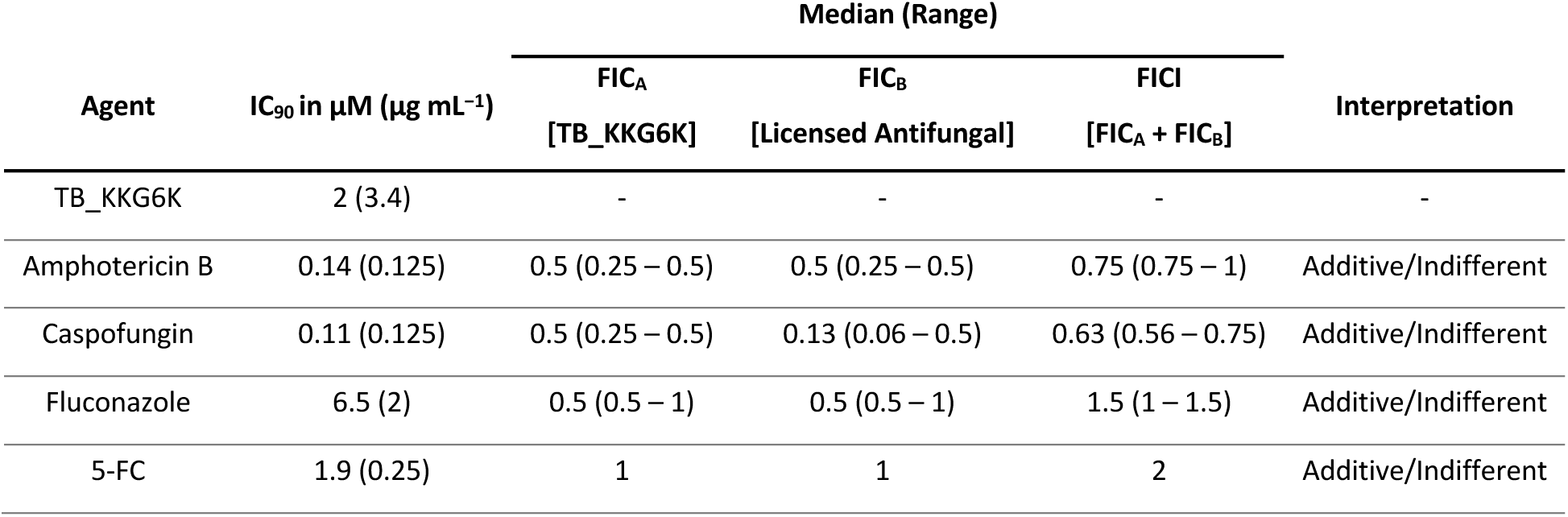
Fractional inhibitory concentration indices (FICI) of licensed drugs and TB_KKG6K against planktonic *C. albicans* cells.

### TB_KKG6K exhibits activity against matured *C. albicans* biofilm

To study the efficacy of TB_KKG6K on sessile *C. albicans* growing on synthetic material, we selected silicone elastomer to first establish the biofilm. Silicone elastomer discs were inoculated with *C. albicans*, after which biofilm development was observed at 24-hour intervals over a 72-hour period using SEM. Following 24 h of incubation, the discs exhibited a moderate degree of biofilm density, comprising predominantly spherical yeast cells (Figure 2A). A 48-hour biofilm exhibited higher cell density, manifesting in the formation of cell clusters as described previously [34, 35]. At this stage of biofilm development, some germinating cells and pseudohyphae, and the formation of ECM between the cells could be observed (Figure 2A, B). Cell density and clustering increased progressively over the next 24 h, ultimately forming a multilayered biofilm by 72 h (Figure 2A) with ECM dense areas (Figure 2C).

**Figure 2.**
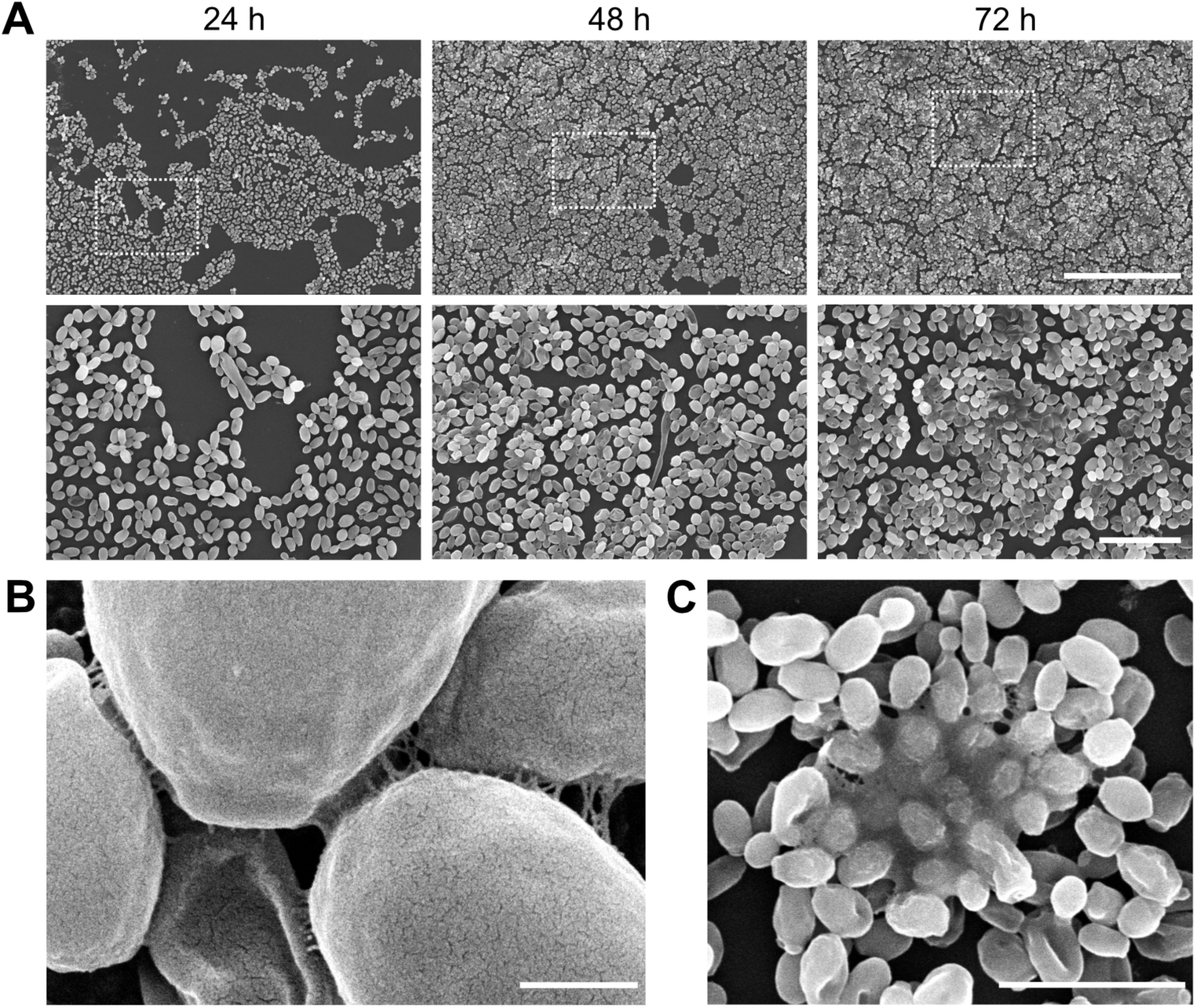
*C. albicans* biofilm and ECM formation. (A) Biofilm development on the silicone elastomer discs was recorded in 24 h-intervals for 72 h. The images were captured with a JSM-6010LV electron microscope at two different magnifications: 400x (upper row), 1,400x (lower row). Scale bars 100 µm and 20 µm, respectively. The dotted square in the images of the upper row indicate the magnified section shown in lower row. (B) A detailed image of ECM formation in a 48 h-old biofilm taken with a TESCAN CLARA electron microscope. Scale bar, 1 µm. (C) Dense accumulation of ECM in a 72 h-old biofilm captured with a JSM-6010LV SEM. Scale bar, 10 µm.

From previous studies, we learned that a concentration higher than the IC_90_, determined through broth microdilution assays on planktonic cells, is required to effectively target *C. albicans* cells growing within a biofilm [20]. Based on this, we applied TB_KKG6K in 25x IC_90_ to evaluate its efficacy on matured biofilms. The treatment of a 48 h-old biofilm with 50 µM TB_KKG6K for 4 h reduced cell clustering, while the cells were severely damaged after a 24 h-treatment in comparison to the untreated biofilm (Figure 3A, C). To capture any adverse effects of the peptide on the yeast cells, we exposed the cells to a subinhibitory concentration (10 µM) of the peptide for the same incubation times (4 h and 24 h). Under these conditions, only few cells with irregular yeast morphology could be observed after 4 h (Figure 3B). The number of these showing a completely collapsed cell structure significantly increased after 24 h (Figure 3D). Similar observations were made in biofilms treated with 10x IC_90_ of amphotericin B (1.25 µg mL^-1^; Figure 3A-D). These effects were absent in the untreated controls (Figure 3B, D).

**Figure 3.**
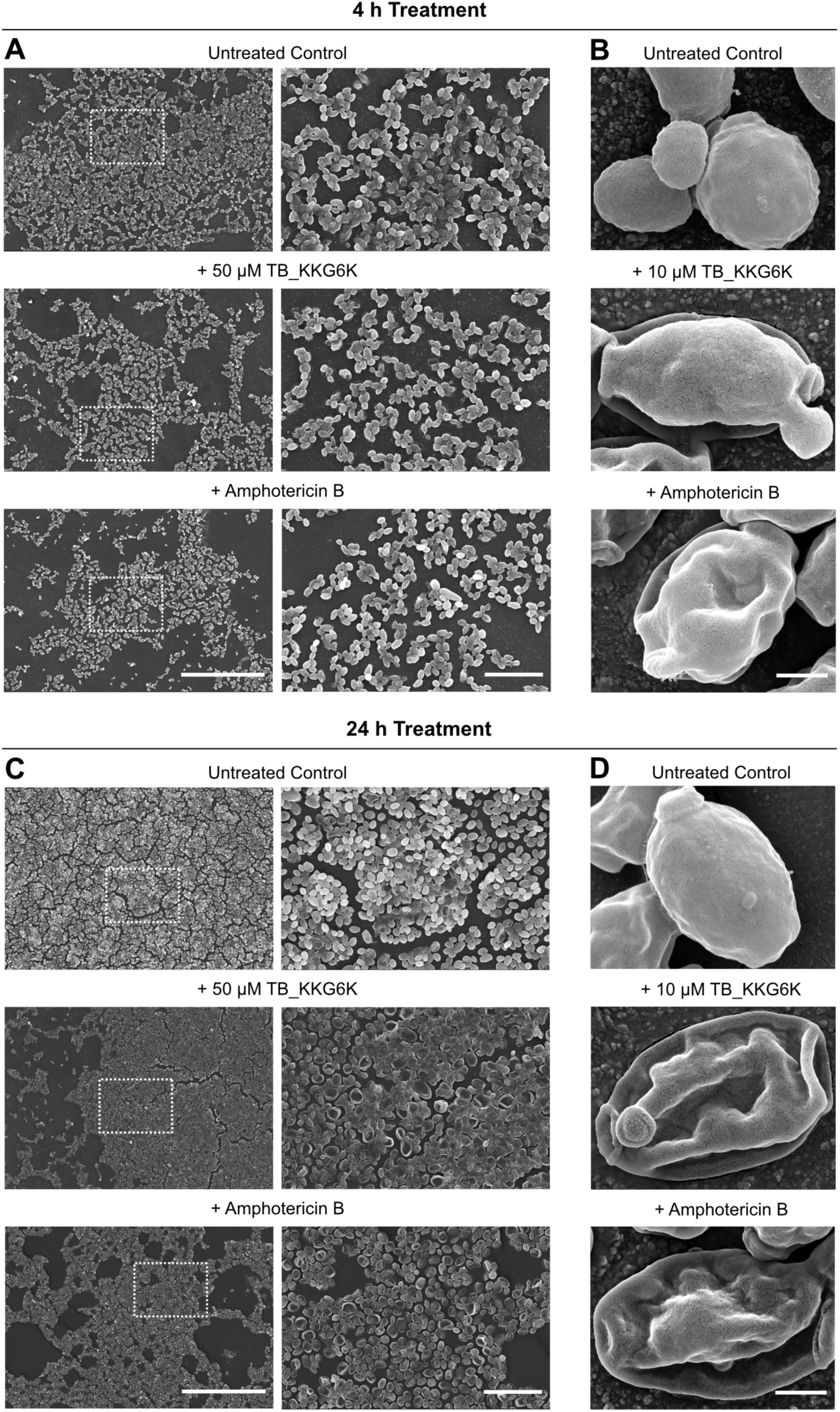
The effect of TB_KKG6K on *C. albicans* cells growing in a biofilm. A 48 h-old biofilm was treated with TB_KKG6K (50 µM) or amphotericin B (1.25 µg mL^-1^) for 4 h (A) or 24 h (C) and compared to the untreated control. The images were captured with a JSM-6010LV electron microscope at two different magnifications: 400x (left), 1,400x (right). Scale bars 100 µm and 20 µm, respectively. The dotted square in the left panels indicate the magnified section shown in the right image. Detailed image of cells grown in a 48 h-biofilm and treated with 10 µM TB_KKG6K or 1.25 µg mL^-1^ amphotericin B for 4 h (B) or 24 h (D). Untreated cells served as control. Images were taken with a TESCAN CLARA SEM. Scale bar, 1 µm.

### TB_KKG6K kills sessile *C. albicans* cells growing in a biofilm

The reduction of sessile *C. albicans* cells embedded in a biofilm was quantified by CFU assay. The number of cells decreased 4 h after the administration of 1x IC_90_ TB_KKG6K, though this change did not reach a level of statistical significance when compared to the untreated control (Table 4). The application of TB_KKG6K at higher concentrations (5-50 µM) evidenced a concentration-dependent reduction, which was statistically significant (Table 4). The administration of 50 µM TB_KKG6K peaked in a decrease of 99.4% of CFUs compared to the untreated control. In a similar manner, amphotericin B treatment reduced the number of CFUs by 99.9%. These reductions correspond to a decrease of viable cells by 2.2 log_10_ and 2.9 log_10_, respectively (Table 4).

**Table 4.**
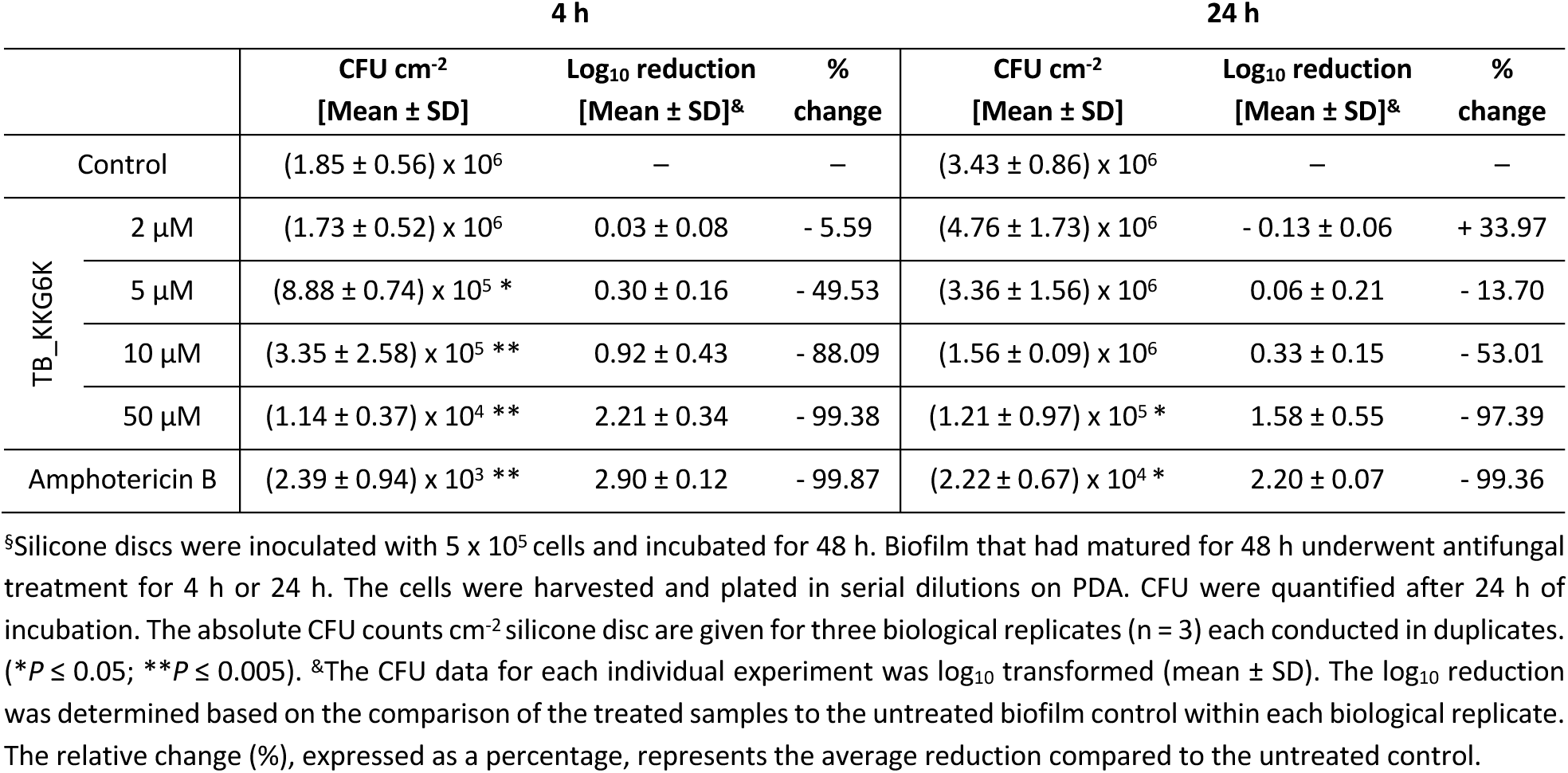
Survival of sessile *C. albicans* cells grown on silicone and treated with increasing concentrations of TB_KKG6K for 4 h or 24 h.^§^.

After 24 h, exposure to peptide concentrations close to the IC_90_ (2-5 µM), did not result in a significant decrease in CFUs compared to the untreated control. Interestingly, there was a slight, though not statistically significant, increase in CFU at 2 µM. However, as the concentration was further increased to 10 µM, a marked decline in the CFU numbers was observed, with the surviving cell population being reduced to 47.0%. At 50 µM TB_KKG6K, this effect was further amplified, leaving only 2.6% of cells viable in the biofilm compared to the untreated control (Table 4). Similarly, treatment with 1.25 µg mL^-1^ amphotericin B resulted in a dramatic reduction of CFUs to 0.6% of surviving cells. Under these test conditions, the antifungal efficacy of TB_KKG6K and amphotericin B against sessile *C. albicans* cells corresponded to a log_10_ reduction of 1.58 and 2.20, respectively (Table 4).

### TB_KKG6K reduces ECM formation in *C. albicans* biofilm

We next analyzed the structural composition of the biofilm using confocal microscopy with the fluorescent stains Calcofluor White and Concanavalin A - AF633 to visualize the cell wall of *Candida* cells and the ECM. In the untreated control, the biofilm exhibited a dense structure predominantly composed of round and oval-shaped yeast cells, including budding, and some germinating and pseudohyphae (Figure 4 A-C). The majority of cells were embedded in ECM, as evidenced by Concanavalin A - AF633 signal, and displayed ECM accumulation in specific regions of the cell surface, reflecting the characteristic heterogeneity of biofilms [7, 36]. The fungal cells were counterstained with Calcofluor White, which binds cell wall chitin. Accumulation of Concanvalin A and Calcofluor White signals at specific sites of the cells coincided also with bud scars [37]. Higher magnification imaging revealed that ECM surrounded the cell wall of *C. albicans*. Treatment with TB_KKG6K resulted in a noticeable reduction in the cell density, accompanied by the presence of collapsed cells (Figure 4C), an effect that paralleled the observations made with amphotericin B treatment and the findings from SEM analysis (Figure 3). Quantification of the total signal intensities of Concanavalin - AF633 revealed a significant reduction in the glucan component of the ECM and cell wall in the TB_KKG6K-treated biofilms compared to the untreated control, similar to the effect observed with amphotericin B (Figure 4D). The Calcofluor White signal, which reflects chitin content, was reduced in the TB_KKG6K-treated biofilms, although the reduction was less pronounced than in the amphotericin B-treated samples, but still notable compared to the untreated control (Figure 4D).

**Figure 4.**
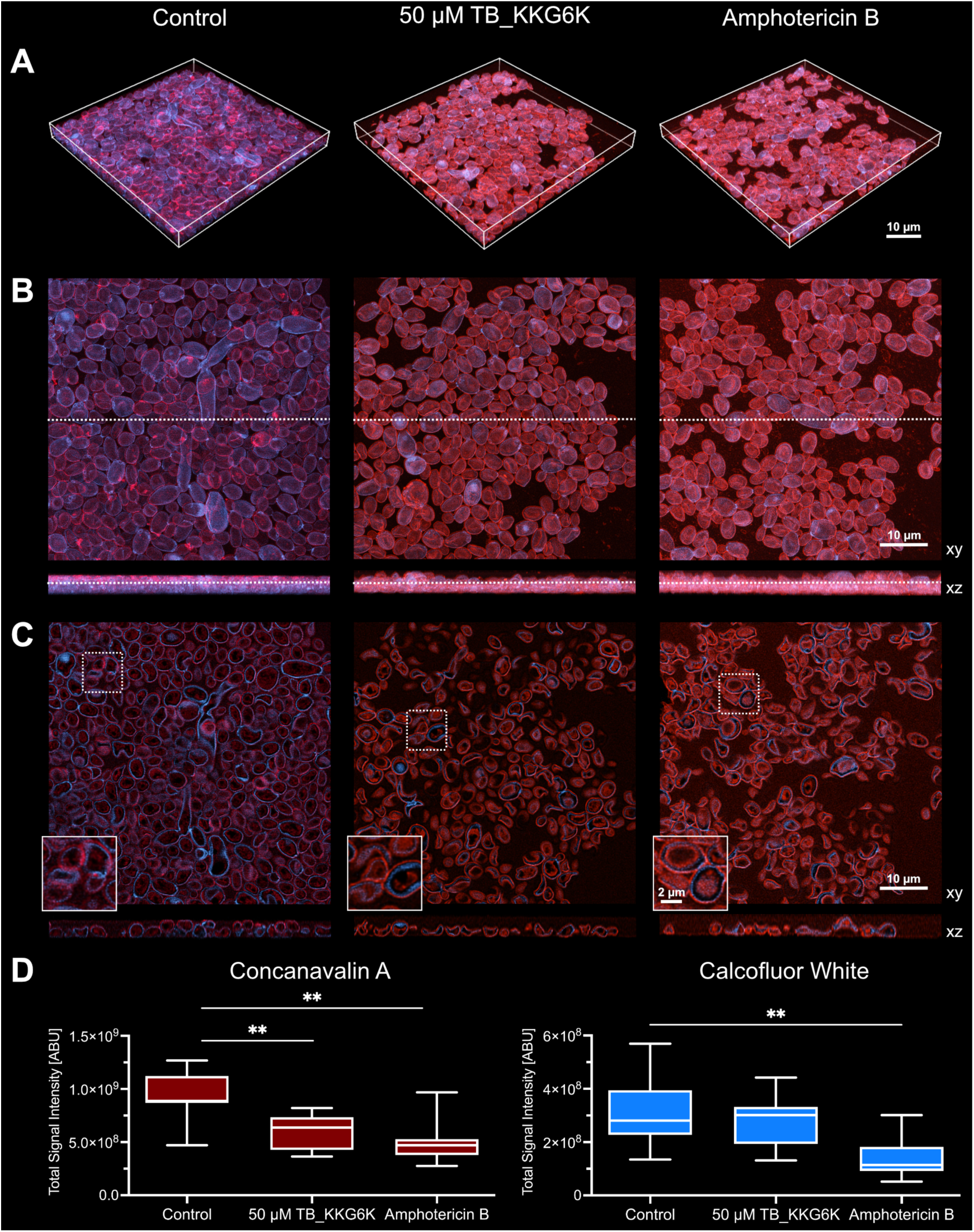
Effect of TB_KKG6K treatment on *C. albicans* biofilm structure. Biofilm that had matured for 48 h was treated with 50 µM TB_KKG6K or 1.25 µg mL^-1^ amphotericin B for 24 h. The fixed samples were stained with 100 µg mL^-1^ Concanavalin A – AF633 and 1 µg mL^-1^ Calcofluor White. Samples were examined using a SP8 gSTED microscope at a magnification of 93x. (A) Images display the maximum intensity profile (MIP) projection in three dimensions (3D). (B) Images present the MIP in two dimensions (2D), with the upper panel oriented in the xy and the lower panel oriented in the xz direction. The dotted lines delineate the region from which sections were selected for depiction in (C). The dotted squares denote the specific regions of xy that were enlarged in the images displayed in the lower left corner of respective images in (C). (D) The total signal quantifications per image was measured for the entire z stack with the Huygens Professional software and is expressed in arbitrary units (ABU). Three representative recordings of each experimental condition from the biological replicates (n = 3) were used, and the resulting data points were arranged in box plots. (\**P* ≤ 0.05 and \*\**P* ≤ 0.005).

### TB_KKG6K induces transcriptional deregulation in sessile *C. albicans* cells

In order to evaluate the nuclear response of *C. albicans* cells growing in a matured biofilm to TB_KKG6K exposure, RT-qPCR was performed. To circumvent global secondary effects induced in dying cells and ensure the recovery of sufficient quantity of material, we applied a sub-lethal peptide concentration (5 µM TB_KKG6K) and a short incubation time (4 h treatment of a 48 h-old biofilm). We investigated the gene transcription of the general biofilm regulating transcription factor *BRG1* and the 1,3-β-D-glucan synthase catalytic subunit *FKS1* using the expression of the house-keeping genes coding for actin (*ACT1*) and elongation factor B (*EFB1*), as previously described [38]. RT-qPCR revealed that TB_KKG6K induced a 2.34-fold increase in the expression of *BRG1* and a 0.64-fold decrease in *FKS1* gene expression (Figure 5).

**Figure 5.**
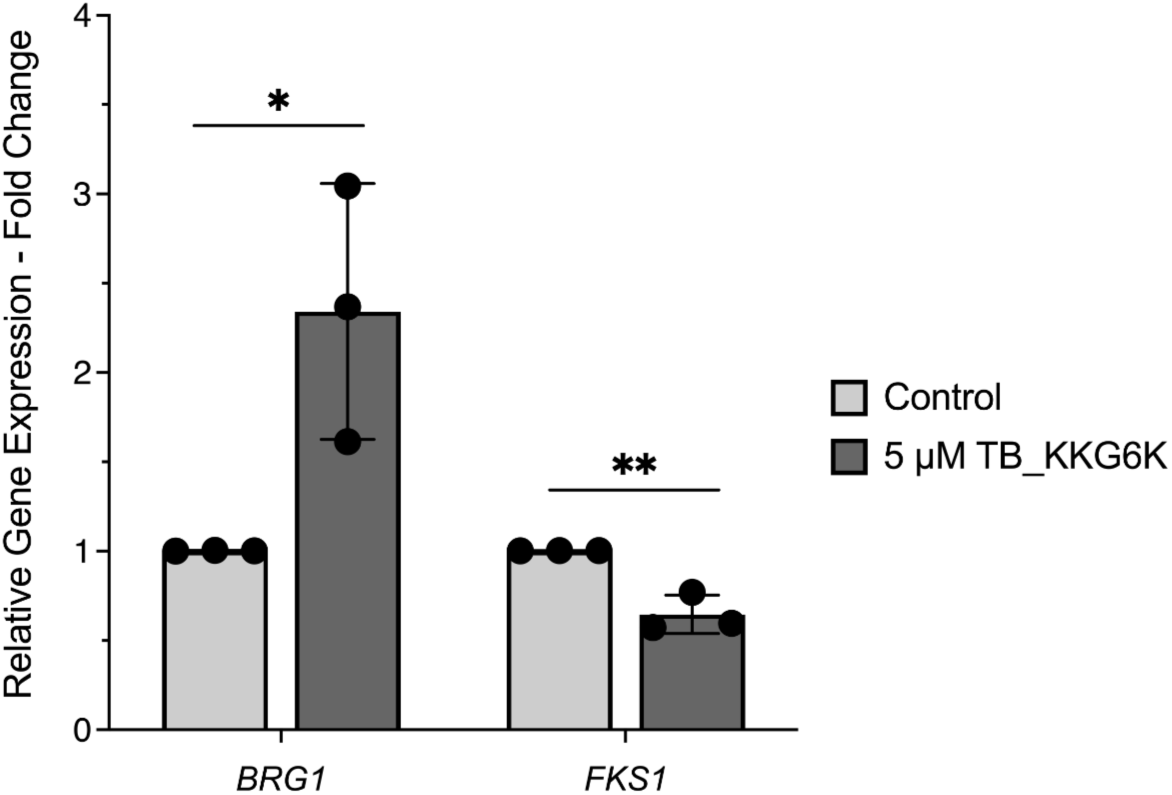
Relative gene expression analysis by RT-qPCR. *C. albicans* biofilm that had matured for 48 h was treated for 4 h with a subinhibitory concentration of TB_KKG6K (5 µM). Gene expression levels were determined using the ΔΔ Ct method. Expression of the target genes *BRG1* and *FKS1* was normalized to the geometric mean of the two housekeeping genes *ACT1* and *EFB1* and is presented as fold change relative to the untreated control. Bars represent mean ± SD from three independent biological replicates (n = 3). Statistical significance was determined by Student’s *t*-test. (\**P* ≤ 0.05 and \*\**P* ≤ 0.005).

## Discussion

This study provides, for the first time, evidence that the amphibian derived TB analog TB_KKG6K has low potential to induce resistance in the opportunistic human pathogenic yeast *C. albicans* and shows the ability to inhibit biofilm maturation on medically relevant material.

*C. albicans* cells showed a mild adaptation to 0.5-1x IC_90_ of TB_KKG6K under prolonged cultivation and were readily killed when transferred to medium supplemented with the peptide at concentrations exceeding 1x IC_90_. This allows the conclusion that the peptide induced tolerance at concentrations ≤ 1x IC_90_, but not the development of resistance mechanisms in planktonic yeast cells. Our observation might be attributable to a complex, potentially multifaceted mode of action of TB_KKG6K. These results align well with our recently published studies, describing the fast fungicidal mode of action in *C. albicans* to be associated with cell membrane activity, entry into the yeast cell, iROS generation, and disintegration of intracellular membranes and organelles [20, 21]. Moreover, TB_KKG6K also inhibited the growth of a fluconazole-resistant *C. albicans* species, which underscore its robust antifungal efficacy [20]. In contrast to TB_KKG6K, however, *C. albicans* was able to survive prolonged subcultivation in the presence of increasing concentrations of the azole drug fluconazole up to 32x IC_90_. Thus, the TB_KKG6K acts differently to fluconazole, which is known to be fungistatic and has a high tendency to induce multiple resistance mechanisms in *C. albicans* [31–33], including the upregulation of drug efflux transporters, mutations in *ERG11*, and overexpression of lanosterol 14-α-demethylase [39–41]. Some AMPs with antifungal activity have been shown to induce tolerance in *C. albicans* upon sequential exposure to increasing peptide concentrations, e.g. fungal NFAP2 and human salivary histatin 3. However, adaptation mechanisms that resulted in tolerance remained unsolved [42, 43]. Future studies need to address whether the *in vitro* results obtained with planktonic cells can be translated to *in vivo* conditions, particularly for cells growing within a biofilm.

Biofilms are defined as complex-surface-associated microbial communities embedded in an ECM composed of glycoproteins, polysaccharides, lipids and nucleic acids. Compared to their planktonic counterparts, biofilm-associated cells show altered gene expression and growth, and are benefitting from ECM mediated protection against environmental stress, immune responses, and antimicrobial agents [44–46]. *C. albicans* biofilms represent a major challenge in the treatment of fungal infections due to their inherent resistance to antifungal drugs. This is particularly problematic, where persistent biofilms lead to severe infections and contamination of medical devices. In one of our previous studies, we could demonstrate that TB_KKG6K exhibited growth-inhibitory efficacy against 24 h-old sessile *C. albicans* cells. However, these biofilms were cultivated on conventional laboratory plastic rather than clinically relevant materials [20]. Therefore, the objective of this study was to characterize in greater detail the activity of TB_KKG6K against matured *C. albicans* biofilms formed on material with clinical relevance.

We selected silicone elastomer, which has found practical application in many biomedical devices, including the tubing of urinary and peritoneal catheters, wound dressing, shunts and drains, contact lenses, orthopedics and many more [47]. Silicone material has been used in studies addressing device-associated *C. albicans* infections [48–50] and natural product-based treatments [51, 52].

TB_KKG6K demonstrated strong efficacy against matured 48 h-old *C. albicans* biofilm on silicone, with a time and concentration dependent reduction in CFU counts determined after treatment within a micromolar peptide concentration range. A statistically significant reduction in biofilm-associated cells, quantified as CFU numbers, was reached after 24 h in the presence of 25x IC_90_ of TB_KKG6K. A similar effect was observed with the drug control sample exposed to 10x IC_90_ amphotericin B. Notably, the decrease in CFU numbers of sessile cells exposed to the peptide for only 4 h was relatively higher when compared to a 24 h treatment. This indicates that cells growing in a biofilm are more vulnerable to the antifungal during a short time exposure than a prolonged exposure. This phenomenon may be explained by a decrease in antifungal efficacy over time. The peptide could be degraded by proteases secreted by the yeast cells, and/or few cells (0.62%) that survived the treatment could resume growth, as quantified by the CFU numbers. These proliferating cells could be a source of secreted protective ECM material. However, technical issues could also explain the above-described phenomenon. Although carefully performed, multiple pipetting steps applied in the preparation of samples for imaging might destabilize cells growing in a biofilm, causing them to detach from the surface and be discharged with the supernatant when it is aspirated off. Therefore, the results obtained through imaging cannot be fully aligned with those obtained through CFU assays. Since antifungal activity in biofilms is both time and concentration dependent, a significant challenge arises in determining whether the drug can be effectively delivered to the site of infection and whether fungicidal concentrations can realistically be achieved and maintained over time. To address this, effective antifungal treatment strategies in the clinics involve the repeated application of a drug to ensure the eradication of the infectious agent.

For analyzing the biofilm structure, the application of SEM was particularly informative in regions of high cell density and ECM accumulation, providing robust evidence for the disruptive potential of the tested peptide against *C. albicans* biofilm integrity. It demonstrated the collapse of *C. albicans* cells grown within the biofilm after treatment with the peptide. Similarly collapsed cells could be observed in the amphotericin B treated control, which was consistent with other studies that showed similar cell deformation to occur in response to drug exposure [53, 54]. No signs of cell wall roughening, pores or other cell surface damages were visible after TB_KKG6K treatment. This aligns with our previous observation obtained with transmission electron microscopy, which revealed the disruption of intracellular membrane structures in planktonic *C. albicans* cells upon peptide treatment, but without pore formation in the cell membrane as an initial trigger for cell death [21].

By using confocal microscopy, we visualized components of the cell wall and the ECM of *C. albicans* cells growing within a biofilm. The cell wall of *C. albicans* exhibits a complex structure, consisting of multiple components organized into two distinct layers. The cell wall’s primary structure is a chitin-β-glucan-mannoprotein framework, wherein chitin is located in the inner layer and glucans together with mannans in the outer layer of the cell wall [55, 56]. Our observations made with confocal microscopy aligned with this structure description (Figure 4). The fluorescent dye Calcofluor White exhibits a high affinity for chitin and cellulose [57]. We observed a mild reduction in Calcofluor White signal intensity in the TB_KKG6K-treated biofilm, while the decrease in signal intensity was more pronounced in biofilms treated with amphotericin B. This might be explained by the ability of this drug to extract ergosterol from the membrane, thereby inducing stress response and impairing the cell membrane function, ultimately compromising the integrity of the inner cell wall [58–62].

The ECM produced by *C. albicans* contains mannans, β-1,6- and β-1,3-glucans, which assemble extracellularly to form a structure that sequesters antifungal agents, blocking their access to cellular targets [63–65]. Concanavalin A, a plant lectin that binds to α-mannopyranosyl and α-glucopyranosyl residues, is used to stain the outer layer of the fungal cell walls and the ECM, when fluorochrom-conjugated [66]. TB_KKG6K, similar to amphotericin B, led to a significant reduction of the Concanavalin A - AF633 signal intensities compared to the untreated controls. This observation can be explained by the reduction of viable cells, documented by a decrease in CFU, and the reduction of cell wall/ECM components through TB_KKG6K. The latter hypothesis is supported by identifying a significant reduction in the expression of the *FKS1* gene encoding the catalytic subunit of the β-1,3-glucan synthase complex with RT-qPCR. Notably, the gene transcription data were collected after a 4 h-long peptide exposure of biofilm, while changes in the cell wall and ECM composition were visualized microscopically after a 24 h-long incubation. However, downregulation of *FKS1* by TB_KKG6K treatment may alter in the long-term cell wall composition and compromise the protective function of ECM, enhancing biofilm disruption. The cell wall and ECM component β-1,3-glucan plays a critical role in the virulence of *C. albicans*. Strains with impaired Fks1 function show reduced fitness, and are less able to undergo yeast-to-hyphae transition and to establish infection *in vivo* [67]. The enzymatic targeting of the ECM with β-1,3-glucanases was shown to increase drug susceptibility and facilitate biofilm eradication [68, 69]. However, based on our previous observation that TB_KKG6K does not localize to the nucleus of the yeast cell [21], we hypothesize that deregulation of gene expression occurs in response to changes in signal transduction, as it has been reported to occur also with other antifungal compounds [70, 71].

*C. albicans* biofilms exhibit a tightly regulated morphogenetic plasticity, characterized by transition between yeast and hyphal forms. A complex signaling network, including master transcription factors such as Brg1, influence the bi-directional transition. These factors control the expression of a coordinated gene network that is essential for dynamic morphological remodeling and biofilm structure [72–74]. SEM and confocal microscopy revealed that *C. albicans* primarily formed a biofilm composed of yeast cells containing only occasionally germinating cells or pseudohyphae, and this observation did not change upon treatment with antifungals. Nevertheless, *BRG1* was found to be overexpressed in the biofilm established on silicone in response to a 4 h TB_KKG6K treatment. We propose that upregulation of *BRG1* could be a compensatory stress response of *C. albicans* aimed at restoring biofilm integrity. Stress-mediated modulation of Tor1 signaling in yeast cells has been reported to be an important factor in the regulation of *BRG1* [29, 75, 76]. This hypothesis certainly merits further analysis in the future.

We could previously show the excellent biocompatibility of TB_KKG6K when applied on keratinocyte-based reconstructed 3D skin models in two independent studies [20, 22]. Its fungicidal mode of action, low risk of resistance development and demonstrated anti-biofilm efficacy, as evidenced in the present study, underscore the significant potential of this peptide for managing chronic or recurrent *C. albicans* infections and preventing colonization of abiotic surfaces. In addition, TB_KKG6K exhibited no antagonistic interactions with standard antifungal drugs, enabling its use in combination therapies at reduced dosages. Notably, TB_KKG6K may be especially suitable for the use in catheter-associated infections, as it is expected to be compatible with silicone materials when delivered in water-based formulations. This makes it a promising candidate for antifungal lock therapy, a treatment approach that involves instilling a high-concentration antifungal agent into colonized intravascular catheters to achieve sterilization, particularly when catheter removal is not feasible and systemic therapy alone is insufficient [77, 78]. An appropriate formulation for the repeated application of TB_KKG6K may be a crucial step to enhance efficacy in anti-*Candida* treatment. Innovative drug delivery approaches offer promising solutions to tackle these challenges, enhancing biofilm penetration and ensuring targeted peptide delivery at the infection site [79]. Furthermore, formulated TB_KKG6K could have great potential to be used in catheter coatings and wound dressings to impede colonization and infection with *C. albicans*, respectively. A feasible example was given with temporin SHa, a member of the temporin family that was successfully grafted onto gold surfaces demonstrating effective surface functionalization [80, 81].

In this study, we utilized a silicone elastomer model that provides a controlled environment for studying biofilm formation and evaluating antifungal treatment on silicone-based devices. However, this model does not account for *in vivo* factors such as host interactions and immune responses and its clinical relevance remains to be determined. Despite these limitations, the study establishes a foundation for future research on treating biofilms on medically relevant materials with natural products like TB_KKG6K.

## Author Contributions

Conceptualization, F.M., C.S. and A.R.; Validation, C.S. and A.G.; Formal Analysis, C.S. and A.G.; Investigation, C.S., A.G., M.K., and A.C.S.; Resources, A.R., M.J.A., D.C.C.-H. and A.C.S.; Writing – Original Draft Preparation, C.S.; Writing – Review & Editing, C.S. and F.M.; Visualization, C.S. and A.G.; Supervision, F.M.; Project Administration, F.M. and C.S.; Funding Acquisition, F.M., C.S. All authors have read and agreed to the published version of the manuscript.

## Acknowledgements

We thank Lena Rieberer for technical assistance, Christopher Spiegel, Martin Offterdinger and Sahana Kale for technical guidance.

## References

1. Nobile, C. J.; Johnson, A. D. *Candida albicans* Biofilms and Human Disease. Annu Rev Microbiol 2015, 69, 71–92. DOI: 10.1146/annurev-micro-091014-104330

2. Gulati, M.; Nobile, C. J. *Candida albicans* biofilms: development, regulation, and molecular mechanisms. Microbes Infect 2016, 18, 310–321. DOI: 10.1016/j.micinf.2016.01.002

3. Atriwal, T.; Azeem, K.; Husain, F. M.; Hussain, A.; Khan, M. N.; Alajmi, M. F.; Abid, M. Mechanistic Understanding of *Candida albicans* Biofilm Formation and Approaches for Its Inhibition. Front Microbiol 2021, 12, 638609. DOI: 10.3389/fmicb.2021.638609

4. Thomas-Rüddel, D. O.; Schlattmann, P.; Pletz, M.; Kurzai, O.; Bloos, F. Risk Factors for Invasive *Candida* Infection in Critically Ill Patients: A Systematic Review and Meta-analysis. Chest 2022, 161, 345–355. DOI: 10.1016/j.chest.2021.08.081

5. Lass-Flörl, C.; Kanj, S. S.; Govender, N. P.; Thompson, G. R.; Ostrosky-Zeichner, L.; Govrins, M. A. Invasive candidiasis. Nat Rev Dis Primers 2024, 10, 20. DOI: 10.1038/s41572-024-00503-3

6. Finkel, J. S.; Mitchell, A. P. Genetic control of *Candida albicans* biofilm development. Nat Rev Microbiol 2011, 9, 109–118. DOI: 10.1038/nrmicro2475

7. Chandra, J.; Kuhn, D. M.; Mukherjee, P. K.; Hoyer, L. L.; McCormick, T.; Ghannoum, M. A. Biofilm formation by the fungal pathogen *Candida albicans*: development, architecture, and drug resistance. J Bacteriol 2001, 183, 5385–5394. DOI: 10.1128/JB.183.18.5385-5394.2001

8. Nett, J.; Andes, D. *Candida albicans* biofilm development, modeling a host-pathogen interaction. Curr Opin Microbiol 2006, 9, 340–345. DOI: 10.1016/j.mib.2006.06.007

9. Malinovská, Z.; Čonková, E.; Váczi, P. Biofilm Formation in Medically Important *Candida* Species. J Fungi (Basel) 2023, 9. DOI: 10.3390/jof9100955

10. d’Enfert, C. Biofilms and their role in the resistance of pathogenic *Candida* to antifungal agents. Curr Drug Targets 2006, 7, 465–470. DOI: 10.2174/138945006776359458

11. Ramage, G.; Martínez, J. P.; López-Ribot, J. L. *Candida* biofilms on implanted biomaterials: a clinically significant problem. FEMS Yeast Res 2006, 6, 979–986. DOI: 10.1111/j.1567-1364.2006.00117.x

12. Talpaert, M. J.; Balfour, A.; Stevens, S.; Baker, M.; Muhlschlegel, F. A.; Gourlay, C. W. *Candida* biofilm formation on voice prostheses. J Med Microbiol 2015, 64, 199–208. DOI: 10.1099/jmm.0.078717-0

13. Bouza, E.; Guinea, J.; Guembe, M. The Role of Antifungals against *Candida* Biofilm in Catheter-Related Candidemia. Antibiotics (Basel) 2014, 4, 1–17. DOI: 10.3390/antibiotics4010001

14. Ponde, N. O.; Lortal, L.; Ramage, G.; Naglik, J. R.; Richardson, J. P. *Candida albicans* biofilms and polymicrobial interactions. Crit Rev Microbiol 2021, 47, 91–111. DOI: 10.1080/1040841X.2020.1843400

15. Mateus, C.; Crow, S. A.; Ahearn, D. G. Adherence of *Candida albicans* to silicone induces immediate enhanced tolerance to fluconazole. Antimicrob Agents Chemother 2004, 48, 3358–3366. DOI: 10.1128/AAC.48.9.3358-3366.2004

16. Mukherjee, P. K.; Chandra, J.; Kuhn, D. M.; Ghannoum, M. A. Mechanism of fluconazole resistance in *Candida albicans* biofilms: phase-specific role of efflux pumps and membrane sterols. Infect Immun 2003, 71, 4333–4340. DOI: 10.1128/IAI.71.8.4333-4340.2003

17. Ramage, G.; Bachmann, S.; Patterson, T. F.; Wickes, B. L.; López-Ribot, J. L. Investigation of multidrug efflux pumps in relation to fluconazole resistance in *Candida albicans* biofilms. J Antimicrob Chemother 2002, 49, 973–980. DOI: 10.1093/jac/dkf049

18. D’Andrea, L. D.; Romanelli, A. Temporins: Multifunctional Peptides from Frog Skin. Int J Mol Sci 2023, 24. DOI: 10.3390/ijms24065426

19. Avitabile, C.; D’Andrea, L. D.; D’Aversa, E.; Milani, R.; Gambari, R.; Romanelli, A. Effect of Acylation on the Antimicrobial Activity of Temporin B Analogues. ChemMedChem 2018, 13, 1549–1554. DOI: 10.1002/cmdc.201800289

20. Kakar, A.; Holzknecht, J.; Dubrac, S.; Gelmi, M. L.; Romanelli, A.; Marx, F. New Perspectives in the Antimicrobial Activity of the Amphibian Temporin B: Peptide Analogs Are Effective Inhibitors of *Candida albicans* Growth. J Fungi (Basel) 2021, 7. DOI: 10.3390/jof7060457

21. Kakar, A.; Sastré-Velásquez, L. E.; Hess, M.; Galgóczy, L.; Papp, C.; Holzknecht, J.; Romanelli, A.; Váradi, G.; Malanovic, N.; Marx, F. The Membrane Activity of the Amphibian Temporin B Peptide Analog TB_KKG6K Sheds Light on the Mechanism That Kills *Candida albicans*. mSphere 2022, 7, e0029022. DOI: 10.1128/msphere.00290-22

22. Schöpf, C.; Knapp, M.; Scheler, J.; Coraça-Huber, D. C.; Romanelli, A.; Ladurner, P.; Seybold, A. C.; Binder, U.; Würzner, R.; Marx, F. The antibacterial activity and therapeutic potential of the amphibian-derived peptide TB_KKG6K. mSphere 2025, 10, e0101624. DOI: 10.1128/msphere.01016-24

23. Meletiadis, J.; Verweij, P. E.; TeDorsthorst, D. T.; Meis, J. F.; Mouton, J. W. Assessing in vitro combinations of antifungal drugs against yeasts and filamentous fungi: comparison of different drug interaction models. Med Mycol 2005, 43, 133–152. DOI: 10.1080/13693780410001731547

24. Kovács, R.; Nagy, F.; Tóth, Z.; Forgács, L.; Tóth, L.; Váradi, G.; Tóth, G. K.; Vadászi, K.; Borman, A. M.; Majoros, L.;, et al. The *Neosartorya fischeri* Antifungal Protein 2 (NFAP2): A New Potential Weapon against Multidrug-Resistant *Candida auris* Biofilms. Int J Mol Sci 2021, 22. DOI: 10.3390/ijms22020771

25. Papp, C.; Kocsis, K.; Tóth, R.; Bodai, L.; Willis, J. R.; Ksiezopolska, E.; Lozoya-Pérez, N. E.; Vágvölgyi, C.; Mora Montes, H.; Gabaldón, T.;, et al. Echinocandin-Induced Microevolution of *Candida parapsilosis* Influences Virulence and Abiotic Stress Tolerance. mSphere 2018, 3. DOI: 10.1128/mSphere.00547-18

26. Geschwindt, A. Studies on the antifungal potential of small, cationic, peptides against *Candida albicans*. Master’s Thesis, Medical University of Innsbruck, Innsbruck, 2023

27. Oliveira, V. C.; Souza, M. T.; Zanotto, E. D.; Watanabe, E.; Coraça-Huber, D. Biofilm Formation and Expression of Virulence Genes of Microorganisms Grown in Contact with a New Bioactive Glass. Pathogens 2020, 9. DOI: 10.3390/pathogens9110927

28. Livak, K. J.; Schmittgen, T. D. Analysis of relative gene expression data using real-time quantitative PCR and the 2(-Delta Delta C(T)) Method. Methods 2001, 25, 402–408. DOI: 10.1006/meth.2001.1262

29. Lu, Y.; Su, C.; Liu, H. A GATA transcription factor recruits Hda1 in response to reduced Tor1 signaling to establish a hyphal chromatin state in *Candida albicans*. PLoS Pathog 2012, 8, e1002663. DOI: 10.1371/journal.ppat.1002663

30. Sah, S. K.; Yadav, A.; Kruppa, M. D.; Rustchenko, E. Identification of 10 genes on *Candida albicans* chromosome 5 that control surface exposure of the immunogenic cell wall epitope β-glucan and cell wall remodeling in caspofungin-adapted mutants. Microbiol Spectr 2023, 11, e0329523. DOI: 10.1128/spectrum.03295-23

31. Sun, L. L.; Li, H.; Yan, T. H.; Fang, T.; Wu, H.; Cao, Y. B.; Lu, H.; Jiang, Y. Y.; Yang, F. Aneuploidy Mediates Rapid Adaptation to a Subinhibitory Amount of Fluconazole in *Candida albicans*. Microbiol Spectr 2023, 11, e0301622. DOI: 10.1128/spectrum.03016-22

32. Gerstein, A. C.; Berman, J. Genetic Background Influences Mean and Heterogeneity of Drug Responses and Genome Stability during Evolution in Fluconazole. mSphere 2020, 5. DOI: 10.1128/mSphere.00480-20

33. Morschhäuser, J. The development of fluconazole resistance in *Candida albicans* - an example of microevolution of a fungal pathogen. J Microbiol 2016, 54, 192–201. DOI: 10.1007/s12275-016-5628-4

34. McCall, A. D.; Pathirana, R. U.; Prabhakar, A.; Cullen, P. J.; Edgerton, M. Author Correction: *Candida albicans* biofilm development is governed by cooperative attachment and adhesion maintenance proteins. NPJ Biofilms Microbiomes 2021, 7, 91. DOI: 10.1038/s41522-021-00264-x

35. Wesenberg-Ward, K. E.; Tyler, B. J.; Sears, J. T. Adhesion and biofilm formation of *Candida albicans* on native and Pluronic-treated polystyrene. Biofilms 2005, 2, 63–71. DOI: doi:10.1017/S1479050505001687

36. Bonhomme, J.; d’Enfert, C. *Candida albicans* biofilms: building a heterogeneous, drug-tolerant environment. Curr Opin Microbiol 2013, 16, 398–403. DOI: 10.1016/j.mib.2013.03.007

37. de Assis, L. J.; Bain, J. M.; Liddle, C.; Leaves, I.; Hacker, C.; Peres da Silva, R.; Yuecel, R.; Bebes, A.; Stead, D.; Childers, D. S.;, et al. Nature of β-1,3-Glucan-Exposing Features on *Candida albicans* Cell Wall and Their Modulation. mBio 2022, 13, e0260522. DOI: 10.1128/mbio.02605-22

38. Hebecker, B.; Vlaic, S.; Conrad, T.; Bauer, M.; Brunke, S.; Kapitan, M.; Linde, J.; Hube, B.; Jacobsen, I. D. Dual-species transcriptional profiling during systemic candidiasis reveals organ-specific host-pathogen interactions. Sci Rep 2016, 6, 36055. DOI: 10.1038/srep36055

39. Whaley, S. G.; Berkow, E. L.; Rybak, J. M.; Nishimoto, A. T.; Barker, K. S.; Rogers, P. D. Azole Antifungal Resistance in *Candida albicans* and Emerging Non-albicans Candida Species. Front Microbiol 2016, 7, 2173. DOI: 10.3389/fmicb.2016.02173

40. Selmecki, A.; Forche, A.; Berman, J. Aneuploidy and isochromosome formation in drug-resistant *Candida albicans*. Science 2006, 313, 367–370. DOI: 10.1126/science.1128242

41. Pristov, K. E.; Ghannoum, M. A. Resistance of *Candida* to azoles and echinocandins worldwide. Clin Microbiol Infect 2019, 25, 792–798. DOI: 10.1016/j.cmi.2019.03.028

42. Fitzgerald, D. H.; Coleman, D. C.; O’Connell, B. C. Binding, internalisation and degradation of histatin 3 in histatin-resistant derivatives of *Candida albicans*. FEMS Microbiol Lett 2003, 220, 247–253. DOI: 10.1016/S0378-1097(03)00121-6

43. Bende, G.; Zsindely, N.; Laczi, K.; Kristóffy, Z.; Papp, C.; Farkas, A.; Tóth, L.; Sáringer, S.; Bodai, L.; Rákhely, G.;, et al. The *Neosartorya (Aspergillus) fischeri* antifungal protein NFAP2 has low potential to trigger resistance development in *Candida albicans* in vitro. Microbiol Spectr 2025, 13, e0127324. DOI: 10.1128/spectrum.01273-24

44. Donlan, R. M. Biofilms: microbial life on surfaces. Emerg Infect Dis 2002, 8, 881–890. DOI: 10.3201/eid0809.020063

45. Flemming, H. C.; van Hullebusch, E. D.; Neu, T. R.; Nielsen, P. H.; Seviour, T.; Stoodley, P.; Wingender, J.; Wuertz, S. The biofilm matrix: multitasking in a shared space. Nat Rev Microbiol 2023, 21, 70–86. DOI: 10.1038/s41579-022-00791-0

46. Lohse, M. B.; Gulati, M.; Johnson, A. D.; Nobile, C. J. Development and regulation of single- and multi-species *Candida albicans* biofilms. Nat Rev Microbiol 2018, 16, 19–31. DOI: 10.1038/nrmicro.2017.107

47. Zare, M.; Ghomi, E. R.; Venkatraman, P. D.; Ramakrishna, S. Silicone-based biomaterials for biomedical applications: Antimicrobial strategies and 3D printing technologies. Journal of Applied Polymer Science 2021, 138, 50969. DOI: 10.1002/app.50969

48. Pierce, C. G.; Chaturvedi, A. K.; Lazzell, A. L.; Powell, A. T.; Saville, S. P.; McHardy, S. F.; Lopez-Ribot, J. L. A novel small molecule inhibitor of *Candida albicans* biofilm formation, filamentation and virulence with low potential for the development of resistance. NPJ Biofilms Microbiomes 2015, 1, 15012-. DOI: 10.1038/npjbiofilms.2015.12

49. Lara, H. H.; Lopez-Ribot, J. L. Inhibition of Mixed Biofilms of *Candida albicans* and Methicillin-Resistant *Staphylococcus aureus* by Positively Charged Silver Nanoparticles and Functionalized Silicone Elastomers. Pathogens 2020, 9. DOI: 10.3390/pathogens9100784

50. McConnell, G.; Rooney, L. M.; Sandison, M. E.; Hoskisson, P. A.; Baxter, K. J. A simple silicone elastomer colonization model highlights complexities of *Candida albicans* and *Staphylococcus aureus* interactions in biofilm formation. J Med Microbiol 2025, 74. DOI: 10.1099/jmm.0.002047

51. Kumpakha, R.; Gordon, D. M. Occidiofungin inhibition of *Candida* biofilm formation on silicone elastomer surface. Microbiol Spectr 2023, 11, e0246023. DOI: 10.1128/spectrum.02460-23

52. Ceresa, C.; Tessarolo, F.; Maniglio, D.; Caola, I.; Nollo, G.; Rinaldi, M.; Letizia, F. Inhibition of *Candida albicans* biofilm by lipopeptide AC7 coated medical-grade silicone in combination with farnesol. AIMS Bioengineering 2018, 5, 192–208. DOI: 10.3934/bioeng.2018.3.192

53. Kim, K. S.; Kim, Y. S.; Han, I.; Kim, M. H.; Jung, M. H.; Park, H. K. Quantitative and qualitative analyses of the cell death process in *Candida albicans* treated by antifungal agents. PLoS One 2011, 6, e28176. DOI: 10.1371/journal.pone.0028176

54. Grela, E.; Zdybicka-Barabas, A.; Pawlikowska-Pawlega, B.; Cytrynska, M.; Wlodarczyk, M.; Grudzinski, W.; Luchowski, R.; Gruszecki, W. I. Modes of the antibiotic activity of amphotericin B against *Candida albicans*. Sci Rep 2019, 9, 17029. DOI: 10.1038/s41598-019-53517-3

55. Gow, N. A.; Hube, B. Importance of the *Candida albicans* cell wall during commensalism and infection. Curr Opin Microbiol 2012, 15, 406–412. DOI: 10.1016/j.mib.2012.04.005

56. Garcia-Rubio, R.; de Oliveira, H. C.; Rivera, J.; Trevijano-Contador, N. The Fungal Cell Wall: *Candida*, *Cryptococcus*, and *Aspergillus* Species. Front Microbiol 2019, 10, 2993. DOI: 10.3389/fmicb.2019.02993

57. Harrington, B. J.; Hageage, G. J., Jr. Calcofluor White: A Review of its Uses and Applications in Clinical Mycology and Parasitology. Laboratory Medicine 2003, 34, 361–367. DOI: 10.1309/eph2tdt8335gh0r3

58. Sokol-Anderson, M. L.; Brajtburg, J.; Medoff, G. Amphotericin B-induced oxidative damage and killing of *Candida albicans*. J Infect Dis 1986, 154, 76–83. DOI: 10.1093/infdis/154.1.76

59. Mesa-Arango, A. C.; Trevijano-Contador, N.; Román, E.; Sánchez-Fresneda, R.; Casas, C.; Herrero, E.; Argüelles, J. C.; Pla, J.; Cuenca-Estrella, M.; Zaragoza, O. The production of reactive oxygen species is a universal action mechanism of Amphotericin B against pathogenic yeasts and contributes to the fungicidal effect of this drug. Antimicrob Agents Chemother 2014, 58, 6627–6638. DOI: 10.1128/AAC.03570-14

60. Mesa-Arango, A. C.; Scorzoni, L.; Zaragoza, O. It only takes one to do many jobs: Amphotericin B as antifungal and immunomodulatory drug. Front Microbiol 2012, 3, 286. DOI: 10.3389/fmicb.2012.00286

61. Anderson, T. M.; Clay, M. C.; Cioffi, A. G.; Diaz, K. A.; Hisao, G. S.; Tuttle, M. D.; Nieuwkoop, A. J.; Comellas, G.; Maryum, N.; Wang, S.;, et al. Amphotericin forms an extramembranous and fungicidal sterol sponge. Nat Chem Biol 2014, 10, 400–406. DOI: 10.1038/nchembio.1496

62. Carolus, H.; Pierson, S.; Lagrou, K.; Van Dijck, P. Amphotericin B and Other Polyenes-Discovery, Clinical Use, Mode of Action and Drug Resistance. J Fungi (Basel) 2020, 6. DOI: 10.3390/jof6040321

63. Dominguez, E.; Zarnowski, R.; Sanchez, H.; Covelli, A. S.; Westler, W. M.; Azadi, P.; Nett, J.; Mitchell, A. P.; Andes, D. R. Conservation and Divergence in the *Candida* Species Biofilm Matrix Mannan-Glucan Complex Structure, Function, and Genetic Control. mBio 2018, 9. DOI: 10.1128/mBio.00451-18

64. Mitchell, K. F.; Zarnowski, R.; Andes, D. R. Fungal Super Glue: The Biofilm Matrix and Its Composition, Assembly, and Functions. PLoS Pathog 2016, 12, e1005828. DOI: 10.1371/journal.ppat.1005828

65. Nett, J.; Lincoln, L.; Marchillo, K.; Massey, R.; Holoyda, K.; Hoff, B.; VanHandel, M.; Andes, D. Putative role of beta-1,3 glucans in *Candida albicans* biofilm resistance. Antimicrob Agents Chemother 2007, 51, 510–520. DOI: 10.1128/AAC.01056-06

66. Tkacz, J. S.; Cybulska, E. B.; Lampen, J. O. Specific staining of wall mannan in yeast cells with fluorescein-conjugated concanavalin A. J Bacteriol 1971, 105, 1–5. DOI: 10.1128/jb.105.1.1-5.1971

67. Ben-Ami, R.; Garcia-Effron, G.; Lewis, R. E.; Gamarra, S.; Leventakos, K.; Perlin, D. S.; Kontoyiannis, D. P. Fitness and virulence costs of *Candida albicans* FKS1 hot spot mutations associated with echinocandin resistance. J Infect Dis 2011, 204, 626–635. DOI: 10.1093/infdis/jir351

68. Tan, Y.; Ma, S.; Leonhard, M.; Moser, D.; Schneider-Stickler, B. β-1,3-glucanase disrupts biofilm formation and increases antifungal susceptibility of *Candida albicans* DAY185. Int J Biol Macromol 2018, 108, 942–946. DOI: 10.1016/j.ijbiomac.2017.11.003

69. Tan, Y.; Ma, S.; Ding, T.; Ludwig, R.; Lee, J.; Xu, J. Enhancing the Antibiofilm Activity of β-1,3-Glucanase-Functionalized Nanoparticles Loaded With Amphotericin B Against *Candida albicans* Biofilm. Front Microbiol 2022, 13, 815091. DOI: 10.3389/fmicb.2022.815091

70. Xie, Y.; Hua, H.; Zhou, P. Magnolol as a potent antifungal agent inhibits *Candida albicans* virulence factors via the PKC and Cek1 MAPK signaling pathways. Front Cell Infect Microbiol 2022, 12, 935322. DOI: 10.3389/fcimb.2022.935322

71. Qian, W.; Lu, J.; Gao, C.; Liu, Q.; Yao, W.; Wang, T.; Wang, X.; Wang, Z. Isobavachalcone exhibits antifungal and antibiofilm effects against *C. albicans* by disrupting cell wall/membrane integrity and inducing apoptosis and autophagy. Front Cell Infect Microbiol 2024, 14, 1336773. DOI: 10.3389/fcimb.2024.1336773

72. Deveau, A.; Hogan, D. A. Linking quorum sensing regulation and biofilm formation by *Candida albicans*. Methods Mol Biol 2011, 692, 219–233. DOI: 10.1007/978-1-60761-971-0_16

73. Cleary, I. A.; Lazzell, A. L.; Monteagudo, C.; Thomas, D. P.; Saville, S. P. BRG1 and NRG1 form a novel feedback circuit regulating *Candida albicans* hypha formation and virulence. Mol Microbiol 2012, 85, 557–573. DOI: 10.1111/j.1365-2958.2012.08127.x

74. Kim, M. J.; Cravener, M.; Solis, N.; Filler, S. G.; Mitchell, A. P. A Brg1-Rme1 circuit in *Candida albicans* hyphal gene regulation. mBio 2024, 15, e0187224. DOI: 10.1128/mbio.01872-24

75. Qi, W.; Acosta-Zaldivar, M.; Flanagan, P. R.; Liu, N. N.; Jani, N.; Fierro, J. F.; Andrés, M. T.; Moran, G. P.; Köhler, J. R. Stress- and metabolic responses of *Candida albicans* require Tor1 kinase N-terminal HEAT repeats. PLoS Pathog 2022, 18, e1010089. DOI: 10.1371/journal.ppat.1010089

76. Su, C.; Lu, Y.; Liu, H. Reduced TOR signaling sustains hyphal development in *Candida albicans* by lowering Hog1 basal activity. Mol Biol Cell 2013, 24, 385–397. DOI: 10.1091/mbc.E12-06-0477

77. Walraven, C. J.; Lee, S. A. Antifungal lock therapy. Antimicrob Agents Chemother 2013, 57, 1–8. DOI: 10.1128/AAC.masthead.57-1

78. Kovács, R.; Majoros, L. Antifungal lock therapy: an eternal promise or an effective alternative therapeutic approach? Lett Appl Microbiol 2022, 74, 851–862. DOI: 10.1111/lam.13653

79. Akkuş-Dağdeviren, Z. B.; Saleh, A.; Schöpf, C.; Truszkowska, M.; Bratschun-Khan, D.; Fürst, A.; Seybold, A.; Offterdinger, M.; Marx, F.; Bernkop-Schnürch, A. Phosphatase-degradable nanoparticles: A game-changing approach for the delivery of antifungal proteins. J Colloid Interface Sci 2023, 646, 290–300. DOI: 10.1016/j.jcis.2023.05.051

80. Lombana, A.; Raja, Z.; Casale, S.; Pradier, C. M.; Foulon, T.; Ladram, A.; Humblot, V. Temporin-SHa peptides grafted on gold surfaces display antibacterial activity. J Pept Sci 2014, 20, 563–569. DOI: 10.1002/psc.2654

81. Oger, P. C.; Piesse, C.; Ladram, A.; Humblot, V. Engineering of Antimicrobial Surfaces by Using Temporin Analogs to Tune the Biocidal/antiadhesive Effect. Molecules 2019, 24. DOI: 10.3390/molecules24040814

